# PARP1/2-linked ADP-ribosylation impairs trophoblast mitochondrial respiration in a TNF-α model of inflammation-mediated preeclampsia

**DOI:** 10.64898/2026.10.20.758860

**Authors:** Abolfazl Nik-Akhtar, Sophia Vanegas Pineda, Jacob Ouellet, Fahmida Jahan, Yusmaris Cariaco, Keir J. Menzies, Shannon A. Bainbridge

## Abstract

Inflammation-mediated preeclampsia (I-PE) is associated with increased placental NAD^+^-dependent ADP-ribosylation and NAD^+^ depletion. We investigated whether PARP1/2- or tankyrase-sensitive ADP-ribosylation contributes to inflammation-associated trophoblast dysfunction and whether pharmacological inhibition improves trophoblast function under inflammatory stress. We profiled mRNA expression of genes encoding enzymes that catalyze poly-ADP-ribosylation (PARP1, PARP2, TNKS1, TNKS2) and selected enzymes that catalyze mono-ADP-ribosylation (*PARP7*, *PARP10*, *PARP12*, and *PARP14*), in human I-PE placentas (control n=23; I-PE n=10) and an LPS-induced rat model of I-PE (control n=8, LPS n=8). HTR-8/SVneo cells were treated with TNF-α alone or with olaparib, (PARP1/2 inhibitor) or XAV939 (TNKS1/2 inhibitor). Global mono- and poly-ADP-ribosylation (MAR/PAR), including a PARP1-associated ∼120-kDa ADP-ribosylation signal and pathway enrichment of proteins identified from this region, were assessed alongside mitochondrial respiration, OXPHOS protein abundance, 4-hydroxynonenal, migration, and viability.

In human I-PE placentas, mRNA abundance of poly-ADP-ribosylating enzymes did not differ significantly from controls, whereas mono-ADP-ribosylating enzymes were increased in LPS-exposed rat placentas. In HTR-8/SVneo cells, TNF-α induced a prominent PARP1 associated ∼120-kDa MAR/PAR signal that was suppressed by olaparib, but not XAV939. Proteins identified from this ∼120-kDa region were enriched for known PARP1 targets and immune/stress-response pathways. Olaparib, but not XAV939, improved basal and maximal respiration, ATP production, and spare respiratory capacity. TNF-α did not significantly alter oxidative phosphorylation (OXPHOS) subunits, 4-HNE abundance, viability, or migration at 24 h.

These data support PARP1/2-sensitive ADP-ribosylation, most consistent with PARP1-linked activity, as a contributor to inflammation-mediated trophoblast bioenergetic dysfunction. The PARP1/2-NAD^+^ axis may therefore represent a mechanism-directed target for future studies of I-PE.

**Graphical abstract:** 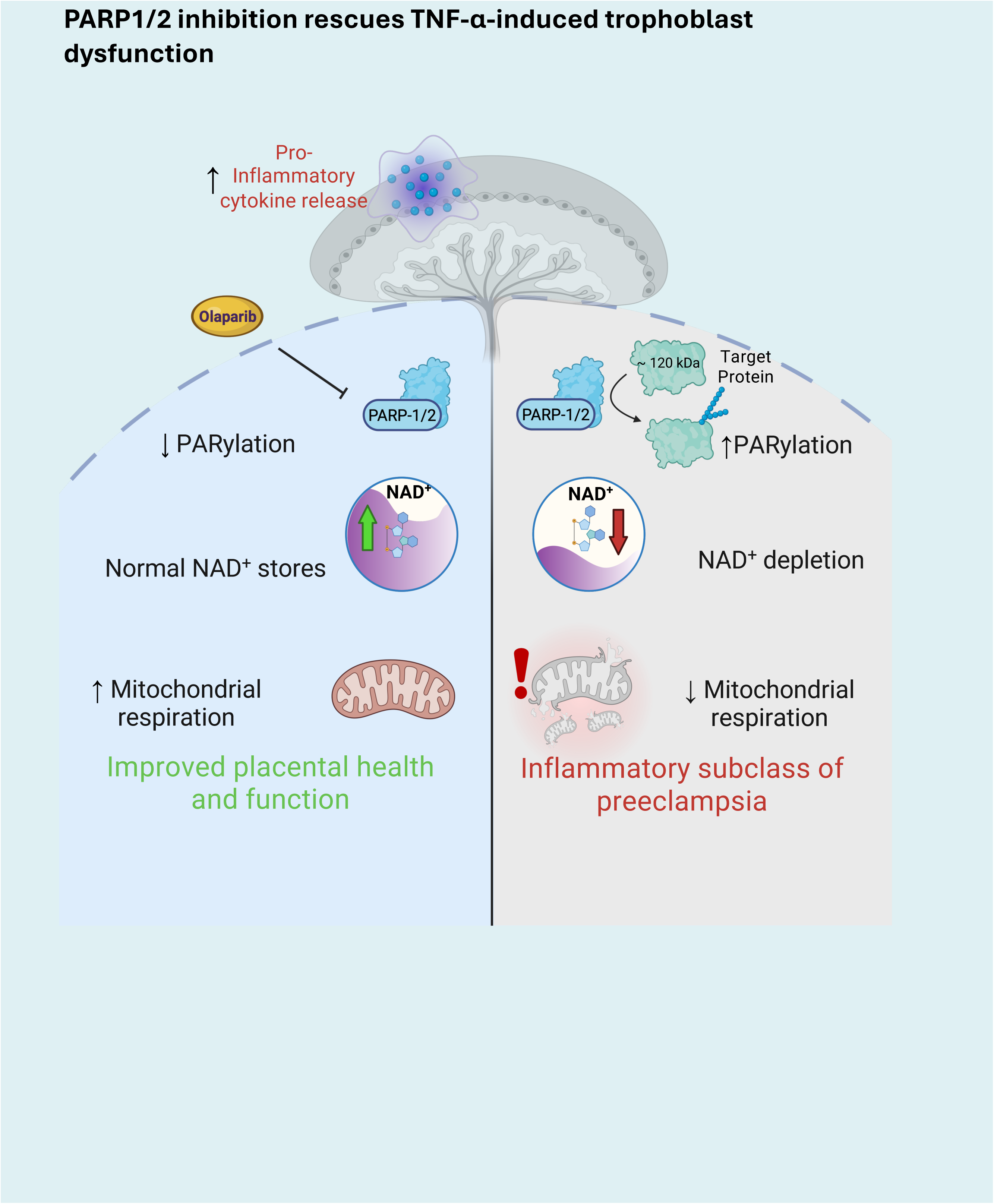

**Highlights:**

- TNF-α-induced PARP1-associated trophoblast ADP-ribosylation (MAR/PAR)
- Olaparib suppresses the TNF-α-induced, PARP1-associated ∼120-kDa MAR/PAR signal
- PARP1/2 inhibition improves TNF-α-impaired mitochondrial respiration
- Tankyrase inhibition does not rescue inflammation-driven trophoblast dysfunction

## Introduction

Preeclampsia (PE) is a major cause of maternal and perinatal morbidity and mortality worldwide, and delivery of the placenta remains the only definitive intervention [1], [2]. The classical two-stage model of PE links impaired spiral artery remodeling to placental malperfusion, hypoxic/oxidative stress, and release of placental stress signals that trigger maternal endothelial dysfunction and clinical disease [3]. However, this framework does not capture the full heterogeneity of PE, as many cases lack defective spiral artery remodeling [4], and molecular, clinical and histological analyses support distinct disease subclasses with divergent etiologies [2], [5], [6].

One clinically relevant subclass is inflammation-mediated PE (I-PE), which is characterized by heightened immune activation at the maternal-fetal interface, as evidenced by the presence of chronic inflammatory placental lesions, and fetal growth restriction despite comparatively milder maternal symptoms [2], [5], [7]. Molecular profiling of I-PE placentas has identified inflammatory cytokine and immune-response signatures, including tumor necrosis factor-α (TNF-α)-associated networks [5], [7], [8]. In this pathological context, TNF-α may contribute to placental dysfunction by impairing trophoblast behavior, barrier function, and vascular interactions [9]. These inflammatory processes may ultimately converge on mitochondrial dysfunction, a feature observed across several PE subclasses [10], [11], [12], [13]. However, the mechanisms directly linking heightened pro-inflammatory signaling to altered trophoblast mitochondrial function and cellular health in I-PE remain incompletely defined.

Our group previously identified a conserved nicotinamide adenine dinucleotide (NAD^+^)-linked injury phenotype across human I-PE placentas, TNF-α-challenged human trophoblast cells, and an LPS-induced rat model of I-PE [14]. Across all three systems, inflammatory stress was associated with increased mono- and poly-ADP-ribosylation (MAR/PARylation)—a major NAD^+^-consuming enzymatic process—leading to depleted NAD^+^ content, impaired mitochondrial function, and oxidative damage. In vitro, these metabolic and stress phenotypes were accompanied by reduced trophoblast viability and invasive capacity, whereas in the rat model of I-PE they were accompanied by altered placental development, reduced placental efficiency, and inflammation-mediated fetal growth restriction. Importantly, NAD^+^ repletion with nicotinamide riboside (NR) attenuated MAR/PARylation, NAD^+^ depletion, mitochondrial dysfunction, and oxidative stress; improved trophoblast viability and invasion in vitro; and improved placental efficiency and fetal growth profiles in vivo [14]. Together, these findings implicate dysregulated MAR/PAR enzyme activity and NAD+ depletion as a candidate mechanistic axis linking inflammatory stress to placental and trophoblast dysfunction in I-PE.

Poly(ADP-Ribose) polymerases (PARPs) belong to the diphtheria toxin-like ADP-ribosyltransferases (ARTDs) family. They are a major class of NAD^+^-dependent ADP-ribosyltransferases that catalyze MAR/PARylation and can influence stress signaling, DNA damage responses, metabolism, and cell function [15], [16], [17], [18]. Excessive PARP activation can deplete NAD^+^ and limit substrate availability for other NAD^+^-dependent pathways, including sirtuins, which regulate mitochondrial function, placental development, and trophoblast differentiation [15], [16]. This inflammation–PARP activation–NAD^+^ depletion–SIRT suppression axis has been linked to mitochondrial dysfunction and tissue injury in non-placental inflammatory diseases (reviewed in [15], [17]). Among the 17 PARP family members, PARP1 and PARP2 are the dominant stress-responsive nuclear ADP-ribosyltransferases, whereas tankyrases (TNKS1/2; PARP5a/5b) are poly-ADP-ribosylating enzymes with distinct roles in Wnt signaling and cell growth [18], [19], [20], [21], [22]. Because broad changes in placental ADP-ribosylation and NAD^+^ depletion have already been demonstrated in I-PE [14], the key unresolved question is which ADP-Ribosylating enzyme activities drive this phenotype and whether selective inhibition of those enzymes could improve trophoblast health under inflammatory conditions.

We therefore profiled candidate ADP enzymes in human I-PE placentas, lipopolysaccharide (LPS)-exposed rat placentas, and TNF-α-challenged human trophoblast cells, and then tested whether selective PARP1/2 or TNKS1/2 inhibition altered inflammatory ADP-ribosylation and trophoblast mitochondrial function in vitro. We hypothesized that TNF-α-induced trophoblast dysfunction would be driven primarily by PARP1/2-sensitive ADP-ribosylation rather than tankyrase activity, and that PARP1/2 inhibition would improve TNF-α impaired mitochondrial respiration.

## Materials and methods

### Human placental samples

Our team previously established a highly annotated archive of placental tissue samples originally obtained from the Research Centre for Women’s and Infants’ Health BioBank, with linked molecular, clinical, and histopathological profiling [5]. For the present study, frozen archived placental biopsies from healthy controls (n = 23) and I-PE cases (n = 10) were selected for qPCR analysis. PE was defined as new-onset hypertension after 20 weeks’ gestation with evidence of maternal end-organ dysfunction and molecular subclassification was performed previously according to published criteria [5]. Human tissue collection and use were approved by the University of Ottawa Research Ethics Board (H08-17-08) and the Mount Sinai Hospital Research Ethics Board (20-0037-E).

### Rodent placental samples

Our team previously developed and characterized an LPS-induced Sprague Dawley rat model of inflammatory PE [14], [23]. Briefly, pregnant rats received escalating intraperitoneal doses of lipopolysaccharide (LPS) from gestational day (GD) 13 to GD18, and placentas were collected on GD19. One placenta was sampled per independent dam/litter. Frozen archived placentas from saline-treated controls (n = 8) and LPS-exposed placentas (n = 8) were used for gene expression analysis in the present study. Animal procedures were approved by the University of Ottawa animal care ethics committee (protocol HS2923) [14], [23].

### HTR-8/SVneo human trophoblast cell culture and treatments

HTR-8/SVneo human trophoblast cells (ATCC CRL-3271; RRID: CVCL_7162; passages 3–18), a model of human extravillous trophoblasts, were cultured in RPMI-1640 (Wisent) supplemented with 10% fetal bovine serum and 1% penicillin/streptomycin at 37°C with 5% CO_2_ and 20% O_2_. To inhibit target enzymes prior to inflammatory challenge, cells were pretreated for 1 h with vehicle (0.1% DMSO), olaparib (3 μM), or XAV939 (10 μM) before stimulation with or without TNF-α (10 ng/mL) for an additional 24 h. Experimental conditions included vehicle control, single treatments (TNF-α, olaparib, or XAV939), dual combinations (TNF-α + olaparib or TNF-α + XAV939), and the triple combination (TNF-α + olaparib + XAV939). TNF-α was selected as an inflammatory stressor consistent with our previous trophoblast model of I-PE [23], and inhibitor conditions were used to compare PARP1/2 sensitive-versus TNKS1/2 -sensitive responses. Unless otherwise indicated, cells were harvested or assayed after 24 h of treatment for RNA, protein, mitochondrial respiration, viability, migration, and oxidative damage analyses. For the early-exposure experiment (Fig. 2C), cells were pretreated with olaparib for 30 min and then exposed to TNF-α for 30 min to assess early ADP-ribosylation dynamics.

### Gene expression analysis

RNA was isolated from human and rat placental tissue and HTR-8/SVneo cells and analyzed by RT-qPCR using SsoAdvanced Universal SYBR Green Supermix (Bio-Rad) and the primer sets listed in Supplementary Table S1. Relative gene expression was calculated using the ΔΔCq method and normalized to the geometric mean of three housekeeping genes (*GAPDH*, *YWHAZ*, and *ACTB* for human samples; *Gapdh*, *Ywhaz*, and *Actb* for rat samples). For HTR-8/SVneo RT-qPCR, 2.5 × 10^5^ cells/well were seeded in six-well plates and allowed to adhere for 24 h. Technical triplicates were used for qPCR; the HTR-8 comparison comprised three independent cell-culture experiments performed on different days.

### Immunoblotting and MAR/PAR signal characterization

Immunoblotting was performed as previously described [24], with slight modifications. Protein lysates were prepared in Pierce RIPA buffer (Thermo Scientific) supplemented with 1 tablet of cOmplete Protease Inhibitor Cocktail (Roche) per 10 ml of RIPA buffer, 1 tablet of PhosSTOP phosphatase inhibitors (Roche) per 10 ml of RIPA buffer, olaparib (Sigma-Aldrich) 10µM, and tannic acid (Sigma-Aldrich) 100µM. Protein concentration was determined using the DC Protein Assay (Bio-Rad), followed by SDS-PAGE and semi-wet transfer to nitrocellulose membranes. Membranes were probed for global mono- and poly (ADP-ribose) (MAR/PAR), OXPHOS complex subunits, and 4-HNE using antibodies listed in Supplementary Table S2. Western blot signals were developed using Clarity Max Western ECL Substrate (Bio-Rad), detected using a ChemiDoc imaging system, and normalized to stain-free total protein. Global MAR/PAR signal, the PARP1-associated ∼120-kDa MAR/PAR signal, PARP1 abundance, OXPHOS complex subunit abundance, and 4-HNE abundance were quantified after 24 h of TNF-α exposure, with or without inhibitor co-treatment. A 30-min TNF-α time point was also assessed to examine early ADP-ribosylation dynamics.

### Proteomic analysis of the ∼120-kDa MAR/PAR-reactive region

To identify proteins within the PARP1-associated ∼120-kDa MAR/PAR region, the corresponding nitrocellulose membrane region was excised and subjected to mass spectrometry-based protein identification using the Orbitrap Astral platform. A same-sized region from the molecular weight marker lane was analyzed as a negative control. Nitrocellulose was dissolved using a modified electroblotted-protein preparation method, substituting acetonitrile (ACN) for acetone. Released proteins were captured using Single-Pot Solid-Phase-enhanced Sample Preparation (SP3): the extract was incubated with 200 μg of a 1:1 Sera-Mag A/B bead mixture for 20 min, washed twice with 100 μL of 80% ethanol and once with 100 μL of 100% ACN, and subjected to overnight on-bead proteolytic digestion in 15 μL of 50 mM ammonium bicarbonate containing protease at 2.5 ng/μL. Digestion was quenched with 5 μL of 0.5% formic acid (FA), and 1 μL was loaded for LC–MS analysis.

Peptides were analyzed using a Vanquish Neo UHPLC coupled to an Orbitrap Astral mass spectrometer (Thermo Fisher Scientific) with positive-mode nano-electrospray ionization. The analytical column (75 μm internal diameter × 150 mm) was packed in-house with 3 μm ReproSil-Pur C18 particles (100 Å; Dr. Maisch). Buffer A was 0.1% FA in water and buffer B was 0.1% FA in 80% ACN. Separation used a 5–40% B gradient over 18.5 min at 500 nL/min. Spray voltage was 2.6 kV and heated-capillary temperature was 300°C. MS and data-independent acquisition (DIA) covered m/z 380–980 with 2-Da DIA windows, 25% HCD energy and MS1 resolution of 240,000. Lock-mass calibration was applied, and data were acquired using Xcalibur.

Raw files were processed in DIA-NN 2.1 in library-free mode against a human UniProt FASTA database. The predicted library used tryptic specificity, one missed cleavage, precursor m/z 380–980 and charge states 2–3, with cysteine carbamidomethylation as a fixed modification and methionine oxidation as a variable modification. Overall and run-specific precursor false-discovery rates were each controlled at 1%. Quantification used QuantUMS with match-between-runs. Identified proteins were then queried against ADPriboDB 2.0 [25] to determine which had prior evidence of MARylation and/or PARylation. Proteins with evidence of PARP1-targeted PARylation were subsequently analyzed by Reactome pathway enrichment analysis.

### Mitochondrial respiration and OXPHOS protein abundance

Mitochondrial oxygen consumption rate (OCR) was measured using the Seahorse XF Cell Mito Stress Test. HTR-8/SVneo cells were seeded at 7.5 × 10^3^ cells/well in XF96 plates, exposed to treatments for 24 h, and assayed in Seahorse XF RPMI medium supplemented with glucose, pyruvate, and glutamine. Oligomycin (1.5 μM), FCCP (1 μM), and rotenone/antimycin A (1 μM) were injected sequentially to derive basal respiration, ATP-linked respiration, proton leak, maximal respiration, non-mitochondrial respiration, spare respiratory capacity, and coupling efficiency. OCR values were normalized to total protein measured by Bio-Rad DC protein assay. Steady-state abundance of representative OXPHOS complex subunits was assessed by immunoblotting, as described above. Basal mitochondrial respiration was calculated as baseline OCR minus non-mitochondrial OCR for each matched experiment.

### Trophoblast viability, migration and oxidative damage assays

Trophoblast migration was assessed with a scratch wound assay [26]. Confluent HTR-8/SVneo monolayers (3 × 10^5^ cells/well in six-well plates) were serum-starved for 5 h, scratched with a sterile pipette tip, and then incubated with the indicated treatment conditions for 24 h. Images were acquired at 0 h and 24 h from the center of each wound. Migration of the HTR-8/SVneo cells into the scratched wound area was quantified as wound confluence at 24 h using an ImageJ-based analysis workflow [27], with greater wound confluence interpreted as increased trophoblast migration/wound closure. Viability was evaluated with calcein-AM (abcam, Cat# ab228556) fluorescence after 24 h of treatment; fluorescence was read at Ex/Em = 485/530 nm using a POLARstar Omega plate reader (BMG Labtech), and H_2_O_2_ (0.8 mM) served as a cytotoxicity control. 4-HNE immunoblotting was assessed as a readout of lipid-peroxidation [6], [28], [29].

### Statistical analysis

Data are presented as mean ± SEM. Two-group qPCR comparisons used Mann–Whitney tests for human placentas (Fig. 1A), unpaired Student’s t-tests for rat PARP/tankyrase expression (Fig. 1B), paired t-tests on log₂-transformed fold-change values for HTR-8/SVneo cells (Fig. 1C), and unpaired Welch’s t-tests on log₂-transformed fold-change values for rat sirtuins (Fig. S1). All tests were two-tailed. Differences among multiple treatment groups were evaluated by ordinary one-way ANOVA for immunoblot endpoints (global MAR/PAR signal, ∼120-kDa PARP1-associated MAR/PAR signal, PARP1 abundance, OXPHOS complex subunits, and 4-HNE), calcein-AM cell viability, and scratch wound confluence. For the Figure 2 immunoblot endpoints, ANOVA was followed by Šídák’s multiple-comparisons test. For the Figure 5 endpoints, ANOVA was followed by uncorrected Fisher’s least significant difference (LSD) tests comparing each treatment with DMSO.

**Figure 1.**
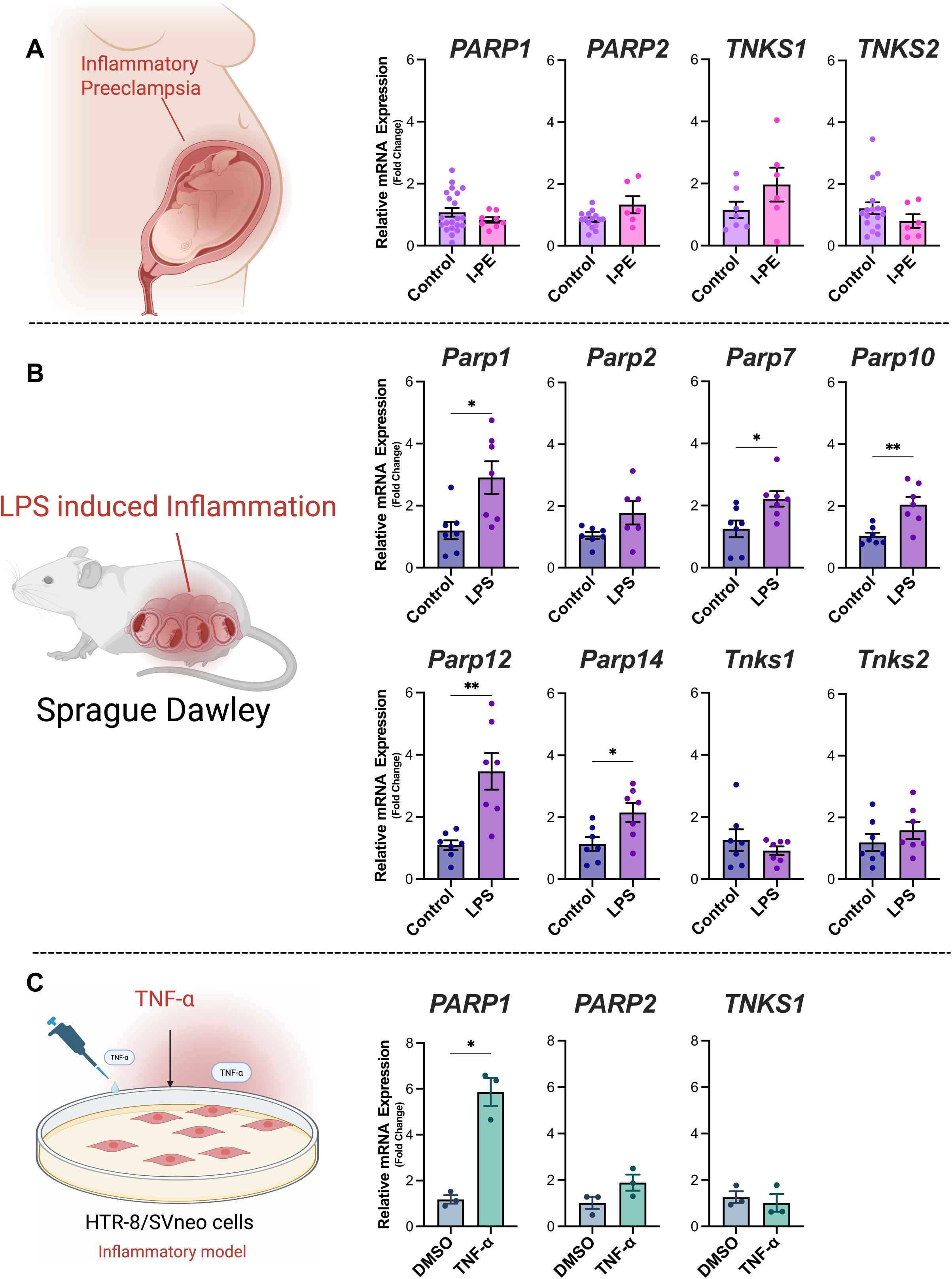
NAD+-consuming enzyme expression across human I-PE placentas and an LPS-induced rat model of I-PE, and TNF-α-treated HTR-8/SVneo cells. (A) Relative mRNA abundance of *PARP1*, *PARP2*, *TNKS1*, and *TNKS2* in human placentas from healthy controls and inflammation-mediated preeclampsia (I-PE) cases. (B) Relative mRNA abundance of *Parp1*, *Parp2*, *Parp7*, *Parp10*, *Parp12*, *Parp14*, *Tnks1*, and *Tnks2* in placentas from saline-treated control rats and LPS-exposed rats. (C) Relative mRNA abundance of *PARP1*, *PARP2*, and *TNKS1* in HTR-8/SVneo cells after 24 h of DMSO or TNF-α treatment in three matched biological experiments. RT-qPCR expression was calculated using the ΔΔCq method with reference-gene normalization and is shown as fold change relative to the corresponding control group. Bars indicate mean ± SEM. Human analyzed sample sizes (control/I-PE) were PARP1, 21/8; PARP2, 15/6; TNKS1, 7/6; and TNKS2, 17/6. Rat analyzed sample sizes were n = 7 per group, except *Parp2* (control n = 7; LPS n = 6). For HTR-8/SVneo cells, n = 3 independent experiments per condition. Human comparisons were analyzed by Mann–Whitney tests, rat comparisons by unpaired Student’s t-tests, and HTR-8/SVneo comparisons by paired t-tests. All tests were two-tailed. *P < 0.05, **P < 0.01 versus the corresponding control.

**Figure 2.**
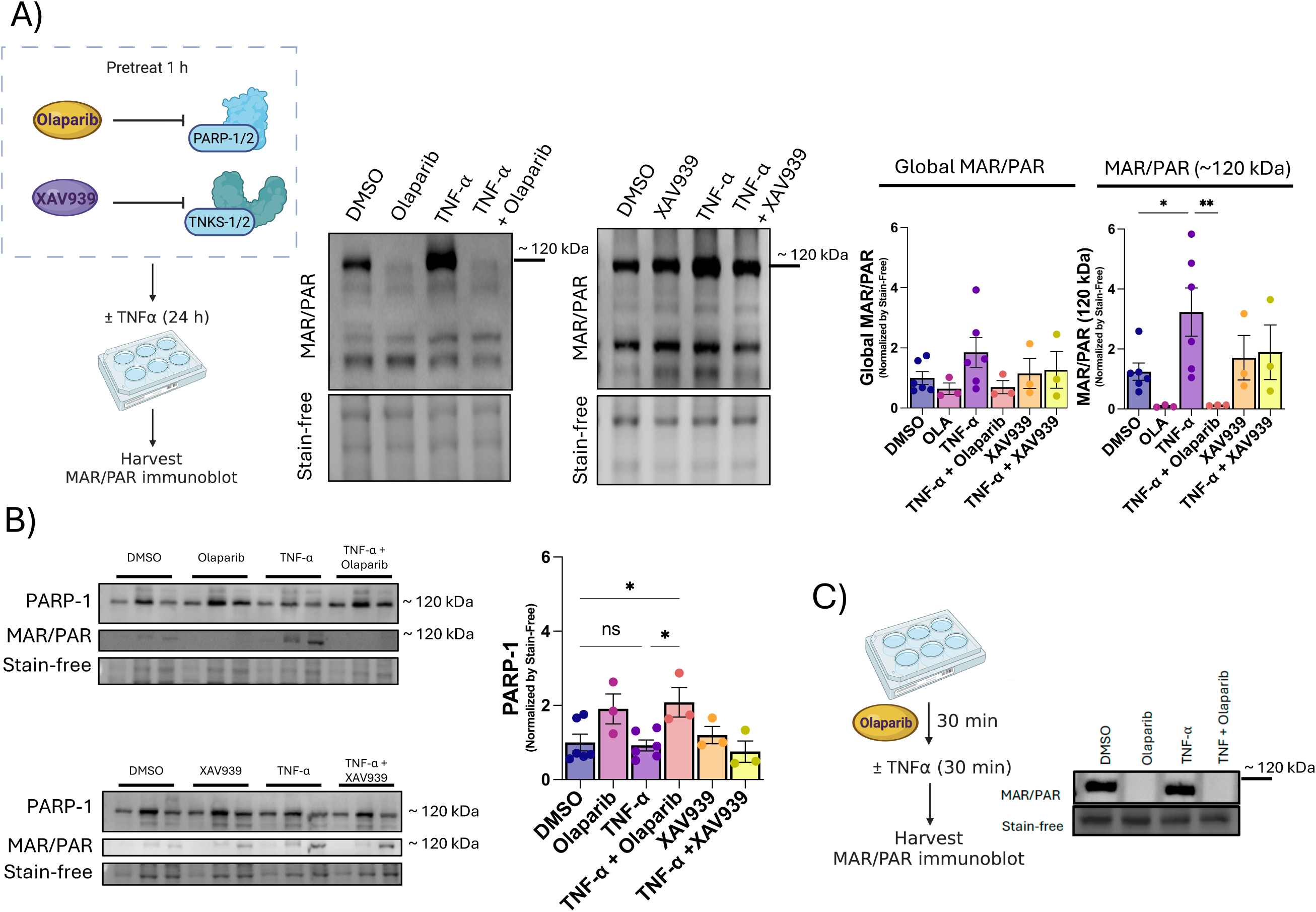
Olaparib, but not XAV939, suppresses TNF-α-induced, PARP1-associated ∼120-kDa ADP-ribosylation signal. HTR-8/SVneo cells were treated with vehicle (0.1% DMSO), TNF-α (10 ng/mL), olaparib (3 μM), XAV939 (10 μM), or the indicated co-treatments. (A) Representative MAR/PAR immunoblots after 24 h of treatment, with densitometric quantification of global MAR/PAR (left graph) and the inducible ∼120-kDa MAR/PAR signal (right graph). (B) Representative PARP1 immunoblots with MAR/PAR re-probing of the same molecular-weight region, and densitometric quantification of PARP1 protein signal. (C) Representative MAR/PAR immunoblot after olaparib pretreatment for 30 min followed by TNF-α exposure for 30 min to assess early ADP-ribosylation dynamics (n = 3 independent experiments). Immunoblot signals were normalized to stain-free total protein. Bars indicate mean ± SEM; points indicate biological replicates. For each quantified endpoint in A and B, n = 6 for DMSO and TNF-α and n = 3 for olaparib, TNF-α + olaparib, XAV939, and TNF-α + XAV939. Quantified data were analyzed by ordinary one-way ANOVA followed by Šídák’s multiple-comparisons test. *P < 0.05, **P < 0.01 for indicated comparisons.

Seahorse OCR-derived parameters were analyzed using repeated-measures one-way ANOVA with Holm–Šídák multiple-comparisons testing, except for proton leak, which was analyzed using a mixed-effects model with Holm–Šídák multiple-comparisons testing to accommodate a single missing value. Normality was assessed using the Shapiro-Wilk test where appropriate, and P ≤ 0.05 was considered statistically significant.

## Results

### Inflammatory placental stress remodels NAD^+^-consuming enzyme expression

To define candidate NAD^+^-consuming enzymes associated with I-PE, we first profiled PARP- and sirtuin-family transcripts across human placental samples and the LPS rat model of I-PE. In archived human I-PE placentas, *PARP1*, *PARP2*, *TNKS1*, and *TNKS2* mRNA abundance did not differ significantly from control placentas (**Fig. 1A**). In contrast, LPS-exposed rat placentas showed increased *Parp1* expression, whereas *Parp2*, *Tnks1*, and *Tnks2* were unchanged (**Fig. 1B**). Several mono-ADP-ribosyltransferase family members, including *Parp7*, *Parp10*, *Parp12*, and *Parp14,* were also increased in LPS-exposed placentas (**Fig. 1B**), together with increased *Sirt1*, *Sirt3*, *Sirt4*, and *Sirt6* expression and decreased *Sirt2* expression (P < 0.05; Fig. S1). These data indicate that inflammatory placental stress is associated with selective remodeling of NAD^+^-consuming enzyme networks, with *Parp1* emerging as the clearest inducible poly-ADP-ribosylating enzyme in the rat I-PE model. Similarly, in HTR-8/SVneo experiments, *PARP1* expression was higher after 24 h TNF-α exposure, whereas *PARP2* and *TNKS1* were unchanged (**Fig. 1C**).

### TNF-α induces an olaparib-sensitive, PARP1-associated ∼120-kDa ADP-ribosylation signal

Because placental ADP-ribosylation is increased in human I-PE [14] and *Parp1* was induced in the rat model of I-PE, we next tested whether inflammatory stress activates a PARP1/2-sensitive ADP-ribosylation response in human trophoblasts cultured in vitro. In HTR-8/SVneo cells, 24 h TNF-α exposure produced only a modest change in global MAR/PAR signal but markedly increased a prominent ∼120-kDa MAR/PAR-positive band (**Fig. 2A**). Olaparib suppressed this ∼120-kDa signal, whereas XAV939 did not, indicating that the TNF-α-induced response is more consistent with PARP1/2-dependent than tankyrase-dependent ADP-ribosylation. Immunoblotting for PARP1 detected the protein in the same molecular-weight region as the inducible MAR/PAR signal (**Fig. 2B**). Given PARP1’s expected molecular weight and known capacity for auto-poly-ADP-ribosylation or poly-ADP-ribosylation by PARP2 [30], the overlapping signals are consistent with ADP-ribosylation of PARP1. However, other ADP-ribosylated proteins of similar apparent molecular weight may also contribute to the MAR/PAR signal. Notably, TNF-α did not increase PARP1 abundance at 24 h, whereas PARP1 signal was higher in olaparib-treated conditions. This pattern suggests that PARP1 protein abundance may be altered in the presence of PARP1/2 inhibition in HTR-8/SVneo, potentially reflecting inhibitor-associated changes in PARP1 stability, turnover, or compensatory regulation instead of inflammatory stimulation alone. Re-probing the same membrane for MAR/PAR again localized the inducible signal to the ∼120-kDa region (**Fig. 2B**). To assess the kinetics of this response, we also examined cells after a shorter 30 min TNF-α exposure. TNF-α rapidly increased the MAR/PAR signal at the ∼120-kDa region, and this early response was blocked by olaparib (**Fig. 2C**). Together, these data support a rapid, olaparib-sensitive, ADP-ribosylation event that co-migrates with PARP1.

### Proteomic analysis links the ∼120-kDa region to PARP1-associated immune/stress pathways

Because the ∼120-kDa region contained the strongest TNF-α-responsive and olaparib-sensitive MAR/PAR signal, and because this signal co-migrated with PARP1 during western blotting but could also include other co-migrating ADP-ribosylated proteins, we next identified proteins present within this molecular-weight window. The corresponding nitrocellulose region was excised and analyzed by LC-MS/MS, followed by comparison with curated ADP-ribosylation and PARP1-target databases (**Fig. 3A**). A total of 267 proteins were identified (including PARP1), of which 203 proteins (76%) had prior evidence of MAR- or PARylation. Among these 203 database-matched ADP-ribosylated proteins, 143 (70%) were annotated as PARP1-associated targets (**Fig. 3B**). Reactome pathway enrichment analysis of the PARP1-associated targets showed strong representation of immune system processes, cytokine signaling, cellular responses to stress, neutrophil degranulation, and broader signal transduction pathways (**Fig. 3C**). Together, these data show that the inducible ∼120-kDa MAR/PAR-reactive region contains numerous proteins with prior evidence of ADP-ribosylation, including many annotated as PARP1-associated targets. Enrichment of immune and stress-response pathways within this protein set further supports a link between this MAR/PAR-reactive region and inflammatory trophoblast signaling.

**Figure 3.**
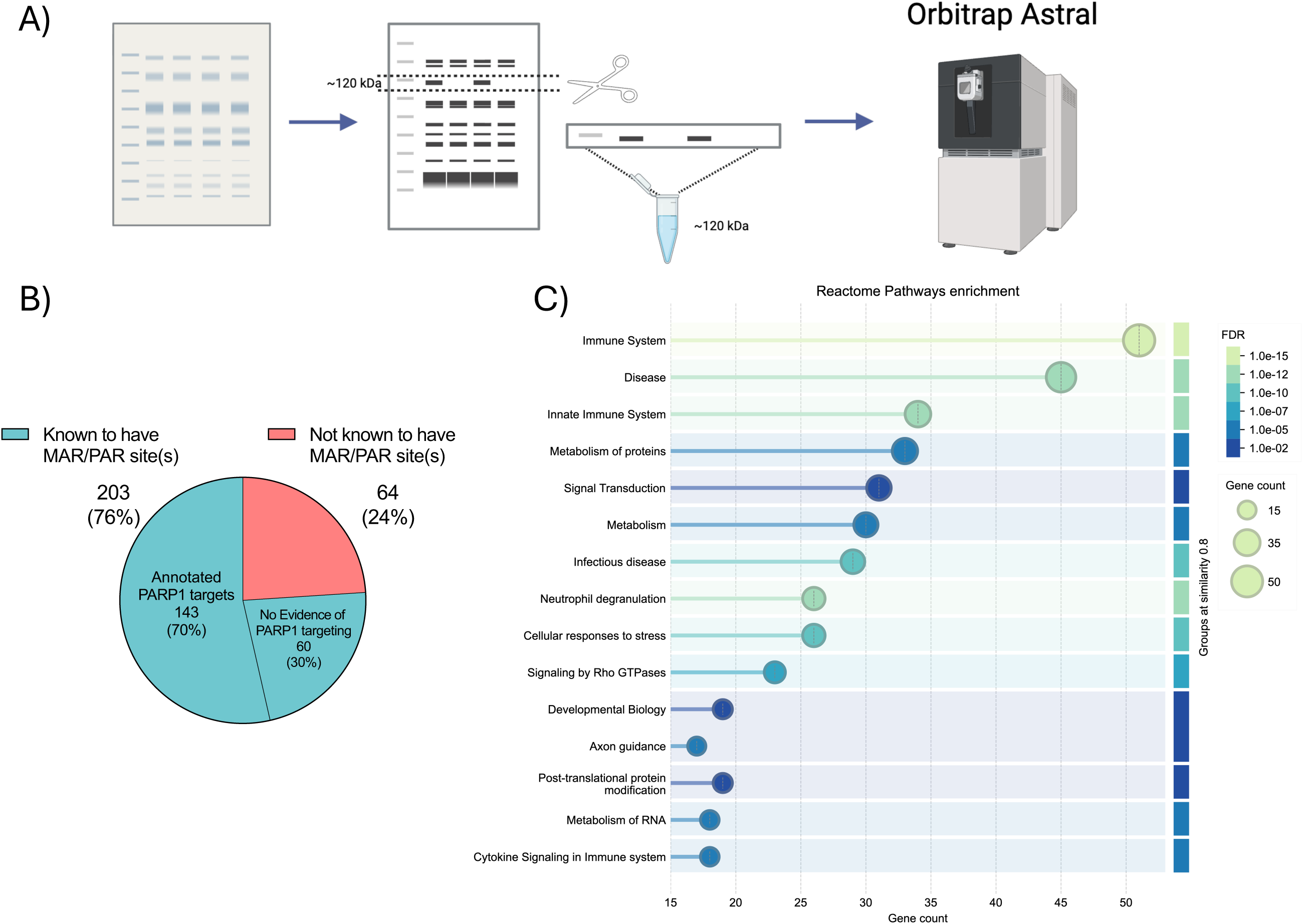
Proteomic profiling links the ∼120-kDa region to PARP1-associated targets and immune/stress pathways. (A) Workflow for excision of the ∼120-kDa MAR/PAR-reactive nitrocellulose region, LC-MS/MS-based protein identification, and downstream annotation using ADP-ribosylation and PARP1-target databases. (B) Proportion of proteins identified within the ∼120-kDa region with prior evidence of MAR/PAR post-translational modification (PTM), and the subset annotated as PARP1-associated targets. Of the 267 proteins identified, 203 (76%) had prior evidence of MAR/PAR modification; within this subset, 143 of 203 (70%) were previously annotated as PARP1-associated targets. (C) Reactome pathway enrichment analysis of PARP1-associated target proteins identified within the ∼120-kDa region.

### PARP1/2 inhibition restores TNF-α-impaired mitochondrial respiration without altering OXPHOS abundance

We next asked whether the olaparib-sensitive ADP-ribosylation response was associated with altered trophoblast bioenergetics. Oxygen consumption rate (OCR) was measured using the Seahorse XF Cell Mito Stress Test, which resolves basal, ATP-linked respiration, proton leak, maximal respiration, spare respiratory capacity, and non-mitochondrial respiration through sequential pharmacological perturbation (**Fig. 4A**). Representative OCR traces showed that TNF-α reduced mitochondrial respiration across the assay, most notably under basal conditions and following FCCP-induced uncoupling, whereas olaparib co-treatment restored the respiratory profile toward control levels (**Fig. 4B**). Quantification confirmed that TNF-α reduced basal respiration and ATP-linked respiration, consistent with impaired steady-state mitochondrial ATP production (**Fig. 4C, D**). TNF-α also reduced maximal respiration and trended to reduce spare respiratory capacity, indicating a diminished ability to respond to increased energetic demand during inflammatory stress (**Fig. 4E, F**). Olaparib co-treatment improved these TNF-α-impaired respiratory parameters, whereas XAV939 did not produce the same rescue in TNF-α-treated cells. Notably, the triple-treatment group, which included TNF-α + olaparib + XAV939, more closely followed the olaparib-rescued phenotype than the TNF-α + XAV939 condition, further supporting PARP1/2-sensitive poly-ADP-ribosylation, rather than tankyrase activity, as the dominant contributor to the TNF-α-induced respiratory defect.

**Figure 4.**
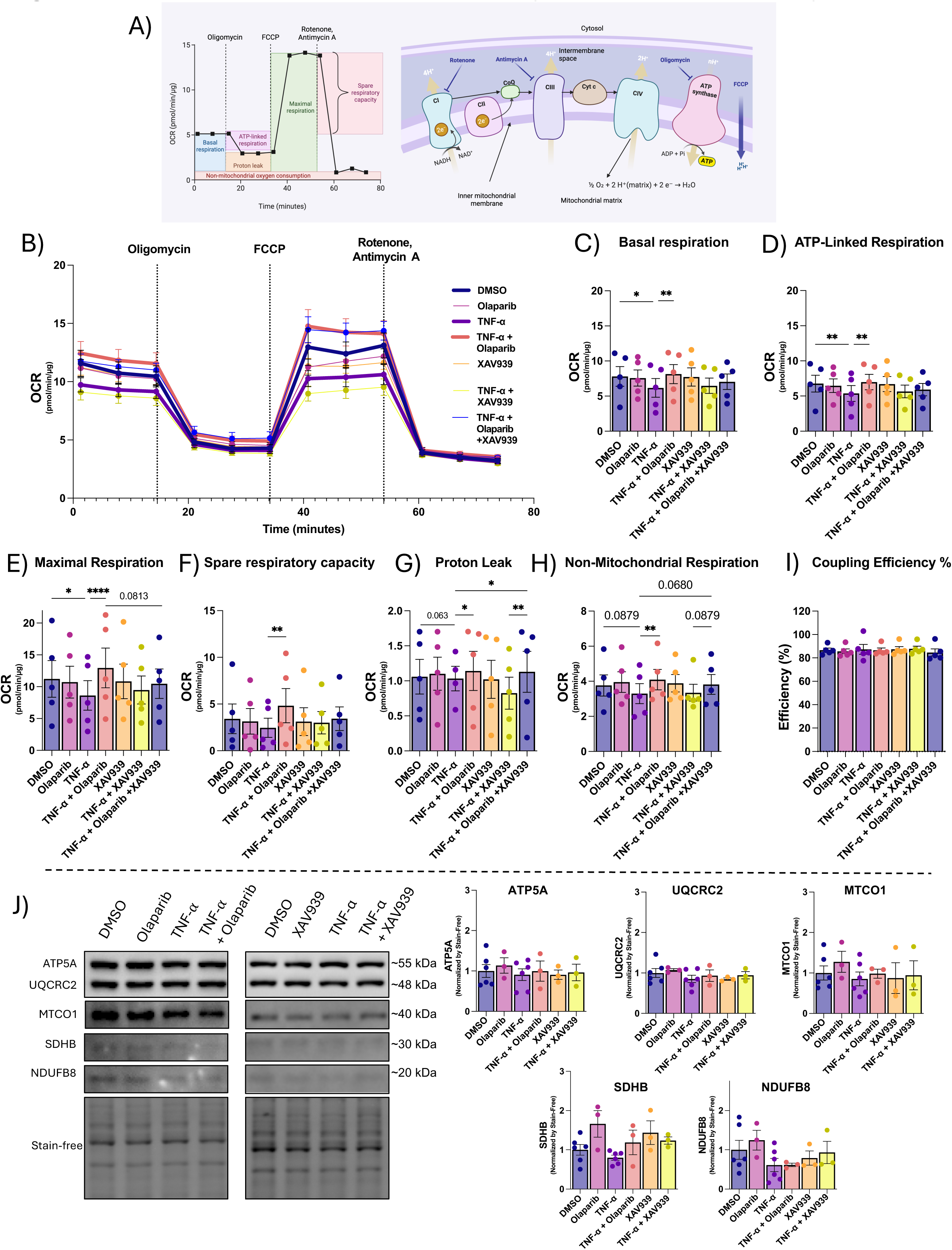
PARP1/2 inhibition restores TNF-α-impaired mitochondrial respiration without major changes in OXPHOS abundance. HTR-8/SVneo cells were treated with vehicle (0.1% DMSO), TNF-α (10 ng/mL), olaparib (3 μM), XAV939 (10 μM), or the indicated co-treatments for 24 h. (A) Schematic overview of the experimental readouts used to assess mitochondrial function. The left schematic shows the Seahorse XF Cell Mito Stress Test workflow, in which oxygen consumption rate (OCR) is measured under basal conditions and after sequential injection of oligomycin, FCCP, and rotenone/antimycin A to resolve ATP-linked respiration, proton leak, maximal respiration, spare respiratory capacity, and non-mitochondrial respiration. The right schematic shows the oxidative phosphorylation (OXPHOS) complexes and representative subunits assessed by immunoblotting: NDUFB8/complex I, SDHB/complex II, UQCRC2/complex III, MTCO1/complex IV, and ATP5A/complex V. (B) OCR traces across the assay. (C–I) Quantification of basal respiration (baseline OCR) (C), ATP-linked respiration (D), maximal respiration (E), spare respiratory capacity (F), proton leak (G), non-mitochondrial respiration (H), and coupling efficiency (I). (J) Representative immunoblots and densitometric quantification of OXPHOS complex subunits. OCR values were normalized to total protein measured by DC assay, and immunoblot signals were normalized to stain-free total protein. In B, lines show mean ± SEM; in C–J, bars indicate mean ± SEM and points indicate biological replicates. For B–I, n = 5 independent experiments per group, except TNF-α in G (n = 4). Panels C, D, E, F, H, and I were analyzed by repeated-measures one-way ANOVA with Holm–Šídák multiple-comparisons testing. Panel G was analyzed by a mixed-effects model with Holm–Šídák multiple-comparisons testing because one value was missing. *\*P < 0.05, **P < 0.01, ***P < 0.001, ****P < 0.0001*.

Proton leak showed only modest treatment-associated changes (**Fig. 4G**). Non-mitochondrial respiration remained within a relatively narrow range, although TNF-α + olaparib showed a small increase compared with TNF-α alone (**Fig. 4H**). Coupling efficiency was not clearly altered across treatments (**Fig. 4I**). Because these Seahorse data indicated a functional respiratory impairment with TNF-α treatment, we next examined the steady-state abundance of representative OXPHOS subunits. Immunoblotting and densitometric analysis of ATP5A, UQCRC2, MTCO1, SDHB, and NDUFB8 did not reveal major treatment-dependent changes (**Fig. 4J**), indicating that TNF-α impairs mitochondrial respiratory function without broadly altering respiratory-chain protein abundance over this time course. This may reflect functional regulation through post-translational modifications, such as ADP-ribosylation.

### TNF-α-induced ADP-ribosylation occurs without overt changes in trophoblast viability, migration, or lipid peroxidation

Finally, we asked whether the TNF-α-induced ADP-ribosylation and respiratory phenotypes were accompanied by overt cytotoxicity, altered trophoblast migration, or lipid peroxidation. Calcein-AM viability assays showed that TNF-α, olaparib, XAV939, or the corresponding co-treatments did not significantly alter cell viability at 24 h, whereas H_2_O_2_ reduced the viability signal as expected (**Fig. 5A**). Because extravillous trophoblast function depends on migratory capacity, we next assessed trophoblast migration using a scratch wound assay. Migration was quantified as wound confluence at 24 h, reflecting the extent of cell movement into the scratched area. TNF-α did not significantly alter wound confluence, and neither olaparib nor XAV939 modified this response (**Fig. 5B**). Finally, 4-HNE immunoblotting showed no treatment-dependent change in lipid peroxidation (**Fig. 5C**). Together, these results indicate that TNF-α induces an olaparib-sensitive ADP-ribosylation response and mitochondrial respiratory impairment without overt loss of viability, migration, or detectable accumulation of 4-HNE at 24 h. This suggests that, under these conditions, the dominant early phenotype is bioenergetic dysfunction but not overt changes in cell function and death.

**Figure 5.**
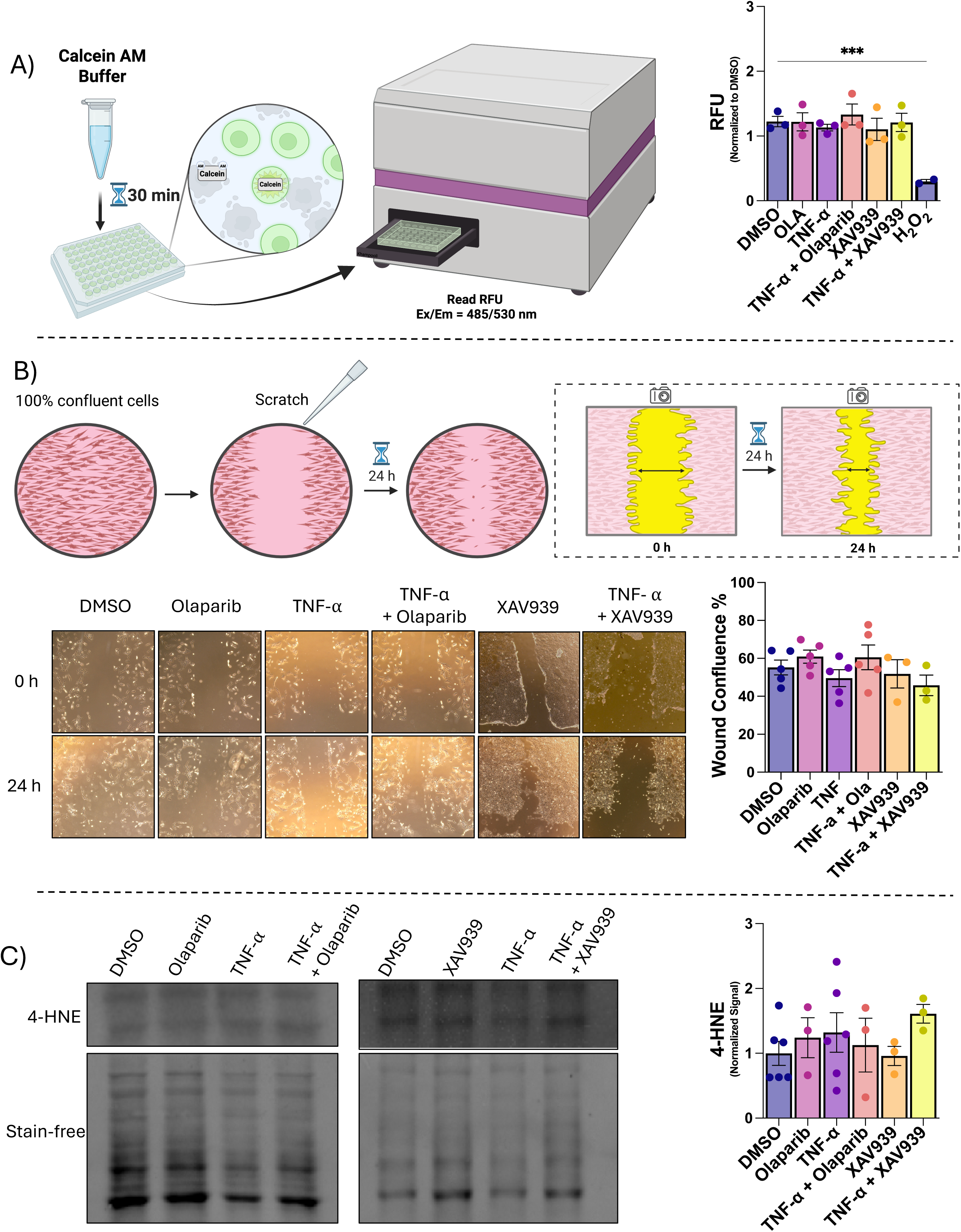
TNF-α-induced ADP-ribosylation occurs without overt changes in trophoblast viability, migration, or lipid peroxidation. HTR-8/SVneo cells were treated with vehicle (0.1% DMSO), TNF-α (10 ng/mL), olaparib (3 μM), XAV939 (10 μM), or the indicated co-treatments for 24 h. (A) Calcein-AM viability assay, with H_2_O_2_ (0.8 mM) included as a positive cytotoxicity control. (B) Scratch-wound assay workflow, representative wound images at 0 h and 24 h, and quantification of wound confluence 24 h after scratching. Greater wound confluence reflects increased cell migration into the scratched area. (C) Representative immunoblots and densitometric quantification of 4-HNE, a marker of lipid peroxidation. Immunoblot signals were normalized to stain-free total protein. Bars indicate mean ± SEM. In A, n = 3 per group, except H₂O₂ (n = 2). In B, n = 5 for DMSO, olaparib, TNF-α, and TNF-α + olaparib and n = 3 for XAV939 and TNF-α + XAV939. In C, n = 6 for DMSO and TNF-α and n = 3 for olaparib, TNF-α + olaparib, XAV939, and TNF-α + XAV939. Panels A, B, and C were analyzed by ordinary one-way ANOVA followed by uncorrected Fisher’s least significant difference (LSD) tests comparing each treatment with DMSO. *P < 0.05, **P < 0.01, ***P < 0.001 for indicated comparisons.

## Discussion

This study was designed to define which poly-ADP-ribosylating enzymes contribute to inflammation-associated trophoblast dysfunction in I-PE and to determine whether selective inhibition of these enzymes improves the trophoblast phenotype. Our previous work established that human I-PE placentas, TNF-α-challenged trophoblasts, and an LPS-induced rat model of I-PE share a conserved NAD^+^-linked injury phenotype characterized by increased ADP-ribosylation, NAD^+^ depletion, mitochondrial dysfunction, and oxidative stress [14]. Here, we extend that work by identifying an olaparib-sensitive, TNF-α-inducible ∼120-kDa MAR/PAR signal that co-migrates with PARP1 immunoreactivity, is enriched for PARP1-associated targets and immune/stress-response pathways and is functionally linked to impaired trophoblast mitochondrial respiration. The lack of a comparable suppression or respiratory rescue with XAV939 indicates that, in this model, the dominant inflammatory ADP-ribosylation phenotype is more consistent with PARP1/2-sensitive activity than tankyrase-dependent signaling. Collectively, these findings support PARP1/2-sensitive ADP-ribosylation as a mechanistic link between inflammatory stress and trophoblast bioenergetic dysfunction in I-PE.

The expression data provide useful context for this mechanism and highlight both model-dependent and stress-responsive regulation of NAD+-consuming enzymes. In the LPS-induced rat model of I-PE, *Parp1* was the clearest inducible poly-ADP-ribosylating enzyme, while several mono-ADP-ribosyltransferases and selected sirtuins were also increased. In TNF-α-treated HTR-8/SVneo trophoblasts, *PARP1* transcripts were increased, indicating that inflammatory stimulation can transcriptionally upregulate this poly-ADP-ribosylating enzyme in this in vitro model. However, the inhibitor data functionally separated these candidates: olaparib suppressed the inducible ∼120-kDa MAR/PAR signal and improved mitochondrial respiration, whereas XAV939 did not. Only *PARP1* showed a consistent increase in TNF-α-treated HTR-8/SVneo cells. The dominant ADP-ribosylation and respiratory phenotypes observed here appear to be driven primarily by PARP1/2-sensitive activity. In contrast, archived human I-PE placentas did not show significant differences in *PARP1*, *PARP2*, *TNKS1*, or *TNKS2* transcript abundance, which may reflect differences between acute inflammatory stimulation and established placental disease, cellular heterogeneity within whole placental biopsies, or RNA-quality constraints associated with archived biobank specimens. Together, these findings indicate that transcript-level changes are informative but do not fully predict the enzyme activity responsible for the phenotype. The rapid increase in MAR/PAR signal within 30 min, together with the absence of TNF-α-induced PARP1 protein accumulation at 24 h, supports a model in which inflammatory stress activates existing PARP1/2-sensitive machinery while also engaging broader transcriptional regulation of NAD+-consuming enzyme networks.

The inducible ∼120-kDa MAR/PAR signal provides the strongest biochemical evidence for a PARP1-associated inflammatory ADP-ribosylation event. PARP1 migrates near this molecular-weight range (∼113-116 kDa) and is known to undergo auto-modification [31], making PARP1 auto-poly-ADP-ribosylation a plausible contributor to the prominent MAR/PAR-reactive band. However, the signal should not be interpreted as definitive evidence that PARP1 is the modified protein in this region. The same molecular-weight window may contain other co-migrating ADP-ribosylated substrates, modified forms of PARP1-associated proteins, or proteins that migrate near PARP1 under the conditions used. This is supported by the proteomic analysis, which identified numerous proteins within the excised ∼120-kDa region, many of which had prior evidence of ADP-ribosylation and were annotated as PARP1-associated targets [32]. Pathway enrichment further linked this protein set to immune signaling, cytokine responses, cellular stress pathways, neutrophil degranulation, and broader signal transduction networks. These findings do not prove that every identified protein is directly ADP-ribosylated in TNF-α-treated trophoblasts, but they do strengthen the interpretation that the dominant MAR/PAR-reactive region represents a PARP1-associated, inflammation-responsive ADP-ribosylation-enriched molecular-weight region rather than a nonspecific global shift in protein modification.

The respiratory data provide the clearest functional link between PARP1/2-sensitive ADP-ribosylation and trophoblast dysfunction. TNF-α reduced basal respiration and ATP-linked respiration, indicating impaired steady-state mitochondrial ATP production [33], and also reduced maximal respiration and spare respiratory capacity, indicating reduced ability to respond to increased energetic demand [34], [35]. Olaparib improved each of these TNF-α-impaired respiratory parameters, whereas XAV939 did not. Moreover, olaparib’s rescue of respiration during TNF-α exposure was largely maintained with or without XAV939, further supporting a contribution of PARP1/2-sensitive activity to the inflammatory respiratory defect. These results fit directly with the NAD^+^-linked mechanism identified previously in I-PE [14], in which inflammatory stress was associated with NAD^+^ depletion and impaired mitochondrial function. Because PARP1/2 activity consumes NAD^+^, excessive PARP1/2-dependent poly-ADP-ribosylation may compromise mitochondrial respiration by limiting NAD^+^ availability for mitochondrial-supportive pathways, including sirtuin-dependent metabolic programs [15], [17], [36].

Importantly, the improvement in mitochondrial respiration occurred without major changes in steady-state OXPHOS complex subunit abundance. TNF-α did not broadly alter ATP5A, UQCRC2, MTCO1, SDHB, or NDUFB8 protein levels over the 24 h treatment period, despite clear suppression of OCR-derived respiratory parameters. This distinction suggests that TNF-α primarily impairs mitochondrial respiratory function rather than causing overt loss of respiratory-chain protein abundance under these conditions. Such a pattern is consistent with a post-translational modification or metabolic mechanism [14], [37], such as altered NAD^+^ utilization, PARP1/2-dependent signaling, or changes in substrate availability, rather than a slower remodeling of mitochondrial protein content. The absence of broad OXPHOS protein changes therefore strengthens rather than weakens the interpretation that PARP1/2-sensitive inflammatory signaling can acutely impair trophoblast bioenergetic performance.

The viability, migration, and lipid peroxidation findings further refine the timing and nature of the phenotype captured in the present study. TNF-α did not significantly reduce calcein-AM viability, impair scratch-wound closure, or increase 4-HNE abundance at 24 h, even though it induced an olaparib-sensitive MAR/PAR signal and impaired mitochondrial respiration. Together, these findings indicate that the 24 h TNF-α exposure used here was sufficient to reveal PARP1/2-sensitive mitochondrial dysfunction, but not sufficient to produce overt cytotoxicity, impaired two-dimensional wound closure, or detectable lipid peroxidation in regular culture media.

This pattern is consistent with the possibility that mitochondrial dysfunction emerges before more severe trophoblast injury in the inflammatory setting, although longer or more complex functional assays are needed to define that progression. Future studies using galactose in place of glucose to increase reliance on oxidative phosphorylation could help determine whether glycolytic compensation limits the functional consequences of TNF-α-induced mitochondrial dysfunction under standard culture conditions. Importantly, these results should not be interpreted as evidence that trophoblast or placental function is broadly preserved during inflammatory stress. In our previous work using the same HTR-8/SVneo model, TNF-α impaired trophoblast invasion through Matrigel, and this defect was rescued by NAD^+^ supplementation [14]. Similarly, our broader work on I-PE showed that inflammatory placental stress is associated with oxidative damage and impaired placental development in vivo, including a reduced exchange region, decreased placental efficiency, and fetal growth restriction [6], [14]. Thus, the absence of altered scratch-wound closure, viability, or 4-HNE accumulation in the present 24 h assay likely reflects the specific functional endpoints and exposure window examined. More complex trophoblast functions, such as matrix invasion, or more chronic inflammatory exposure may be required for bioenergetic dysfunction to progress into overt cellular and placental functional deficits.

The lack of effect of XAV939 across the major biochemical and functional endpoints is also mechanistically informative. Tankyrases are poly-ADP-ribosylating enzymes and have established roles in Wnt signaling, cell growth, and stress-associated pathways [18], [38], but XAV939 did not suppress the TNF-α-induced ∼120-kDa MAR/PAR signal or rescue mitochondrial respiration. These findings argue that tankyrase inhibition is neither sufficient nor necessary to explain the inflammatory ADP-ribosylation and respiratory phenotype observed here. This does not exclude tankyrase-dependent biology in other placental contexts or at other time points, but it narrows the dominant enzyme-sensitive signal in this TNF-α trophoblast model toward PARP1/2-sensitive activity.

This study has limitations that should be considered when interpreting the findings. Although HTR-8/SVneo cells provide a tractable human trophoblast model, they do not recapitulate the cellular complexity, oxygen gradients, immune interactions, or endocrine environment of the inflamed placenta. A second consideration is that the human qPCR analysis was performed using archived placental material, which may have reduced sensitivity to detect modest transcript-level differences. The proteomics analysis also has interpretive constraints: proteins identified within the ∼120-kDa MAR/PAR-reactive region were annotated using existing ADP-ribosylation and PARP1-target databases [25], [32] and these data should therefore be interpreted as enrichment of a PARP1-associated protein set rather than direct substrate mapping. Finally, olaparib was used here as a mechanistic probe of PARP1/2-sensitive activity and should not be interpreted as a pregnancy-ready therapeutic candidate, particularly given the embryo-fetal toxicity observed with systemic olaparib in animal studies [39]. Nevertheless, the olaparib-sensitive rescue of mitochondrial respiration remains important because it implicates PARP1/2-dependent NAD^+^ consumption as a modifiable candidate mechanism linking inflammatory stress to trophoblast bioenergetic dysfunction. This distinction supports future work aimed at defining pregnancy-compatible strategies to preserve NAD^+^ availability, limit maladaptive ADP-ribosylation, or restore mitochondrial function in I-PE.

Overall, this work advances the I-PE mechanism from a broad NAD+-linked injury phenotype to a more defined, enzyme-sensitive pathway connecting inflammation to trophoblast bioenergetic dysfunction. Rather than reflecting generalized ADP-ribosylation or overt cellular injury, the TNF-α response was marked by a selective olaparib-sensitive, PARP1-associated ∼120-kDa MAR/PAR signal and a respiratory defect that emerged without detectable changes in OXPHOS protein abundance, viability, migration, or lipid peroxidation at 24 h. This distinction is important because it positions PARP1/2-sensitive ADP-ribosylation as an early and functionally relevant stress response, capable of compromising mitochondrial performance before more overt indices of trophoblast injury are apparent. By separating PARP1/2-sensitive effects from tankyrase inhibition, these findings refine the enzymatic source of the NAD+-linked phenotype in I-PE and identify trophoblast mitochondrial respiration as a key functional consequence of inflammatory PARP1/2 activation.

## Acknowledgments

The authors acknowledge the technical support provided by the University of Ottawa proteomics core.

## Author contributions (CRediT)

**Abolfazl Nik-Akhtar:** Investigation, Formal analysis, Visualization, Writing – review & editing. **Sophia Vanegas Pineda:** Conceptualization, Investigation, Formal analysis, Visualization, Writing – original draft. **Jacob Ouellet:** Formal analysis, Visualization. **Fahmida Jahan:** Investigation, Formal analysis, Visualization, Writing – review & editing. **Yusmaris Cariaco:** Formal analysis, Visualization, Writing – review & editing. **Keir J. Menzies:** Conceptualization, Methodology, Resources, Supervision, Funding acquisition, Writing – review & editing. **Shannon A. Bainbridge:** Conceptualization, Methodology, Resources, Supervision, Funding acquisition, Writing – review & editing.

## Ethics approval and informed consent

Human tissue use was approved by the University of Ottawa Research Ethics Board (protocol H08-17-08) and the Mount Sinai Hospital Research Ethics Board (protocol 20-0037-E). The parent rodent study was approved by the University of Ottawa animal care ethics committee (protocol HS2923). Human participants provided informed consent for placental tissue banking and research use.

## Data availability

Data supporting the findings of this study are available from the corresponding authors upon request.

---

**Supplementary Figure S1.**
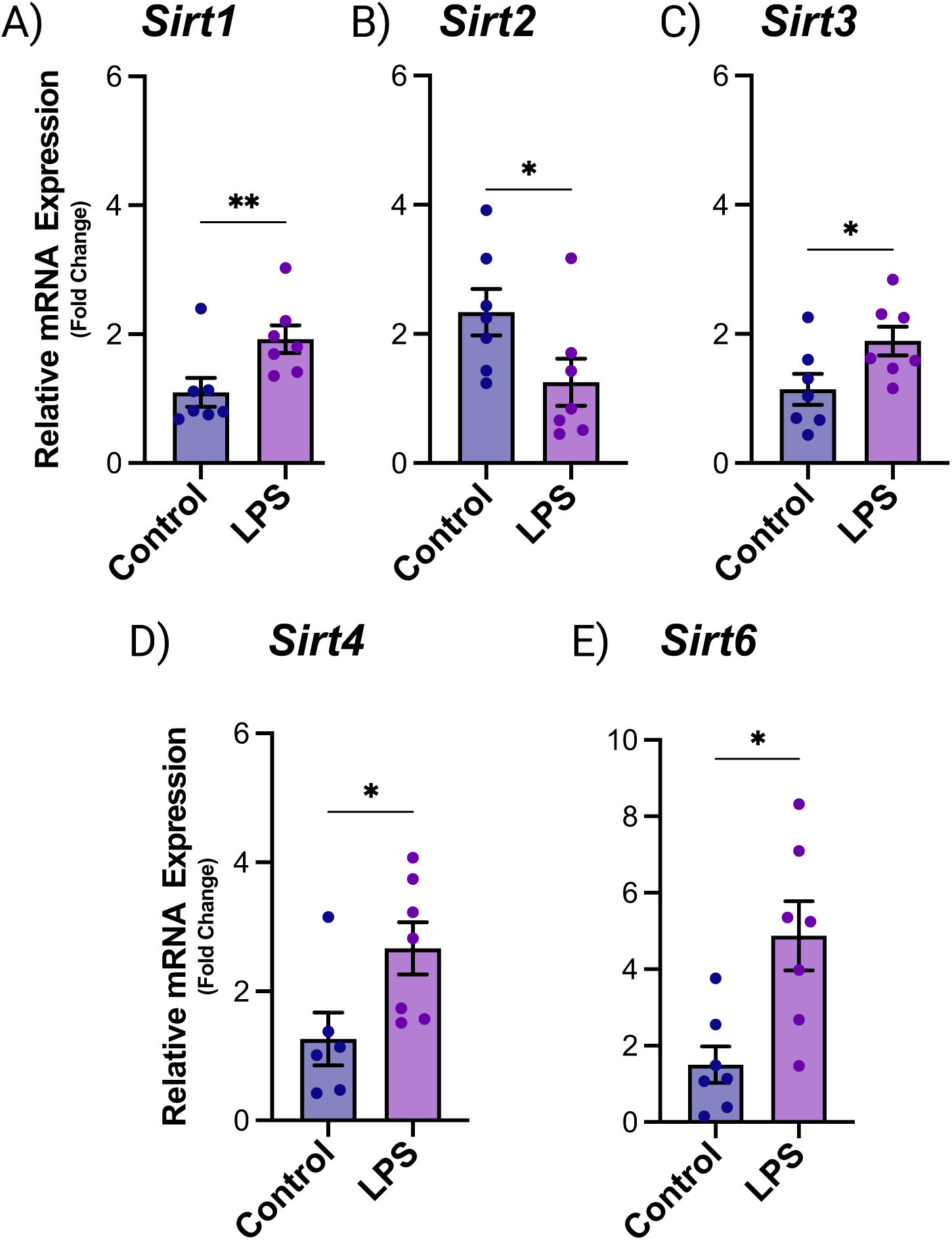
Sirtuin transcript expression in placentas from an LPS-induced rat model of inflammation-mediated preeclampsia. Relative mRNA abundance of *Sirt1* (A), *Sirt2* (B), *Sirt3* (C), *Sirt4* (D), and *Sirt6* (E) in placentas from saline-treated control rats and LPS-exposed rats. Expression was measured by RT-qPCR using the ΔΔCq method, normalized to the geometric mean of *Gapdh*, *Ywhaz*, and *Actb*, and expressed as fold change relative to the control group. Bars indicate mean ± SEM of fold-change values; each point represents one placenta from an independent dam/litter. In A, B, C, and E, n = 7 per group; in D, control n = 6 and LPS n = 7. All five genes were analyzed using unpaired two-tailed Welch t-tests on log₂-transformed fold-change values. *P < 0.05, **P < 0.01 versus control.

