## Supplemental Tables for "PARP1/2-linked ADP-ribosylation impairs trophoblast mitochondrial respiration in a TNF-α model of inflammation-mediated preeclampsia"

### Supplementary tables

Supplementary Table S1. Primer pairs for RT-qPCR.

| Species / role | Gene | Forward primer (5' → 3') | Reverse primer (5' → 3') |
| --- | --- | --- | --- |
| Human target | *PARP1* | TGGAAAAGTCCCACACTGGTA | AAGCTCAGAGAACCCATCCAC |
| Human target | *PARP2* | GGCACAAATCAAGGCAGGTTA | AAGTCATGCGGAATCCTGGTG |
| Human target | *TNKS1* | TGGATCTCTGGCAGTTTACTCC | TCAATCTCTCCCTAAGCTCCG |
| Human target | *TNKS2* | GGGTGTCCAGTTCACAAAGA | AAGCCTGTTCTCCTCTGTAAATAA |
| Rat target | *Parp1* | TTGACTATGGCCAGGACGAAGA | CTGCCTTCTGCTCAGTTTCCC |
| Rat target | *Parp2* | CCAAGGTCGGAAAGGCTCAT | TGTGCTGCCCTGTTTTCCCA |
| Rat target | *Parp7* | ACTTTGATCCCCGTGTCTGT | GCCCATTGTGTATCTGCCAG |
| Rat target | *Parp10* | ATGACTGGCTTTCGGCTCTG | GCACAACACATGGCAGCTAA |
| Rat target | *Parp12* | GCCATGACTTACGGTGCTACC | CCAAACTCATCACTCCAGTACCA |
| Rat target | *Parp14* | CACCTGGAAGATGATGGAGCCA | GAGGTTCACTTTCTGCTGCACC |
| Rat target | *Tnks1* | ACGGCTCTGACCCTTCCATC | GGCTGCACAGTTGCTTCACA |
| Rat target | *Tnks2* | TAGCAGACCCATCCGCCAAA | TTCTCCTCATTGCCACTCCTGG |
| Rat target | *Sirt1* | AAGTTGACTGTGAAGCTGTACG | TGCTACTGGTCTTACTTTGAGGG |
| Rat target | *Sirt2* | CACGCAGAACATAGATACCCTG | CAGTGTGATGTGTAGAAGGTGC |
| Rat target | *Sirt3* | CCCCAAGCCCTTTTTCACTTT | CGACACTCTCTCAAGCCCA |
| Rat target | *Sirt4* | CCGTAGAGCTGTGAGAGAATGA | ACCCAATGGAGGCTTTCGAG |
| Rat target | *Sirt6* | TGTTTGTGGAAGAATGTGCCA | CCTTAGCCACGGTGCAGAG |
| Human reference | *ACTB* | GCACAGAGCCTCGCCTTT | TATCATCATCCATGGTGAGCTGG |
| Human reference | *GAPDH* | AGCCACATCGCTCAGACAC | GCCCAATACGACCAAATCC |
| Human reference | *YWHAZ* | TGTAGGAGCCCGTAGGTCATC | GTGAAGCATTGGGGATCAAGA |
| Rat reference | *Actb* | ATTGGCAATGAGCGGTTC | GGATGCCACAGGACTCCAT |
| Rat reference | *Gapdh* | GTGAAGGTCGGTGTGAACGG | TGAACTTGCCGTGGGTAGAGT |
| Rat reference | *Ywhaz* | GGTCATCTTGGAGGGTCGTCT | CTGCGAAGCATTGGGGATCAA |

**Supplementary Table S2. Primary antibodies.**

| Target | Manufacturer | Catalog number or clone |
| --- | --- | --- |
| Poly/Mono-ADP-ribose | Cell Signaling Technology | E6F6A |
| PARP1 | Santa Cruz Biotechnology | sc-8007 |
| OXPHOS cocktail | Abcam | ab110413 |
| 4-Hydroxynonenal (4-HNE) | R&D Systems | MAB3249 |
